# Whales and Men: genetic inferences uncover a detailed history of hunting in bowhead whale

**DOI:** 10.1101/2020.04.09.033191

**Authors:** TB Hoareau

**Affiliations:** Reneco International Wildlife Consultants LLC, Sky Tower, office 3902 and 3903 - Al Reem Island P.O. Box 61741 - Abu Dhabi, United Arab Emirates; Institution at which research was done: Molecular Ecology and Evolution Programme, Department of Biochemistry, Genetics and Microbiology, University of Pretoria, Pretoria 0002, South Africa

**Keywords:** Abundance, *Balaena mysticetus*, Extended Bayesian Skyline plot, Last Glacial Maximum, Population dynamics, Whaling

## Abstract

After millennia of hunting and a population collapse, it is still challenging to understand the genetic consequences of whaling on the circumarctic bowhead whale. Here I use published modern mtDNA sequences from the Bering-Chukchi-Beaufort population and a new time calibration to show that late–glacial climate changes and whaling have been the major drivers of population change. Cultures that hunted in the Arctic Seas from as early as 5000 years ago appear to be responsible for successive declines of the population growth, bringing the effective size down to 38% of its pristine population size. The Thules and the Basques (year 1000–1730) who only hunted in the North Atlantic had a major impact on this North Pacific population, indicating that bowhead whale stocks respond to harvesting as a single population unit. Recent positive growth is inferred only after the end of commercial whaling in 1915, and for levels of harvesting that are close to the current annual quota of 67 whales. By unfolding the population history of the bowhead whale, I provide compelling evidence that mtDNA yields critical yet undervalued information for management and conservation of natural populations.

## Introduction

For over four millennia, man has been hunting bowhead whales in the circumarctic Seas (Savelle and Kishigami 2013; Seersholm et al. 2016). Basque whalers recorded a consequential population decline in the late 1500s (Barkham 1984), and by 300 years later, most populations had collapsed (Ellis 1991). The vulnerability of bowhead whales to exploitation and climate change (Foote et al. 2013) is exacerbated by a slow population turn-over (George et al. 2011) and a disappearing sea-ice habitat (Lindsay and Schweiger 2015; Chambault et al. 2018). Nonetheless, populations have been increasing since the end of commercial whaling in 1915 (Witting 2013), and despite multiple attempts, the previous severe decline remains undetected using genetics (Rooney et al. 1999 and 2001; Borge et al. 2007; Givens et al. 2010; Phillips et al. 2013).

Genetic bottlenecks can be challenging to detect, even in well documented demographic collapses (Peery et al. 2012). However, researchers using coalescent-based full Bayesian methods (e.g. Heled and Drummond 2008) have shown promising results (Fontaine et al. 2012; Trucchi et al. 2016; Mays et al. 2018), especially when used in tandem with ancient DNA (Lorenzen et al. 2011; Beland et al. 2019). The success of these methods on modern datasets partly depends on the use of appropriate time calibrations, yet despite recent efforts (Crandall et al. 2012; Hoareau 2016), adequate mutation rates are still scarce, even for the bowhead whale.

With the cumulative effects of human activities on habitat and climate, providing a detailed population history for wild species becomes urgent for improved conservation management, especially when bottlenecks and low genetic diversity can compromise adaptive evolutionary potential and cause extinction (Frankham et al. 1999; Ørsted et al. 2019). To reconstruct the population history of the bowhead whale and unravel the potential impact of known historical drivers, I reanalysed 2494 bp of mtDNA sequence for 149 specimens (Phillips et al. 2013) from the Bering-Chukchi-Beaufort region (BCB). This population is showing signs of recovery, has far greater abundance than others (Moore et al. 2019), and has been monitored and studied for several decades (Zeh and Punt 2005). After inferring female effective population size *(Ne)* using a Bayesian skyline analysis (Heled and Drummond 2008), I calculated population growth rates; a parameter that has a key role in describing population trends and identifying potential drivers of decline (Sibly and Hone 2002). I then applied a molecular clock derived from a single-point calibration and used a clustering method that groups the population growth rates into distinct time periods. Remarkably, I found that bowhead whale demography is strongly influenced by sea-level changes and five millennia of human harvesting.

## Results and discussion

### Validation of an LGM-based calibration of the molecular clock

To calibrate the molecular clock, I assumed an initial population expansion co-occurring with the onset of the Northern hemisphere deglaciation at 20 ka (Clark et al. 2009), and applied the hypothesis that subsequently, the abundance of the whale has increased with its core suitable habitat (Foote et al. 2013). Simulations show that such a Last Glacial Maximum (LGM) point–calibration is pertinent for species that experienced a demographic expansion associated with deglaciation (Supplementary Fig. 1). Applied to the bowhead whale mtDNA dataset, the calibration provided a mutation rate of 12.3×10^−8^ per bp per year (Supplementary Fig. 2). This is close to rates obtained for bowhead whale using ancient DNA (15.9×10^-8^ per bp per year; Ho et al. 2008), but 10-fold higher than the rate derived from fossil calibration (e.g. 1.2× 10^-8^ per bp per year; Phillips et al. 2013) that biases time and population parameters (Ho et al. 2011). The substantial variation in population growth rates over the last 20 kyr consistently groups into nine demographic periods (Supplementary Fig. 3–4) when applying a Bayesian clustering method (Fraley and Raftery 2003). The transition times estimated between these periods (Supplementary Fig. 5 and Methods) help us compare genetically inferred demographic changes and potential environmental and anthropogenic drivers that are known to have influenced the abundance of the whale.

### Effect of climate change during the late glacial period

The first demographic period, at 20 ka, can be inferred to have started with an effective population size of 4510 individuals (*Neτ*. effective population size scaled to the generation time *τ*) (Supplementary Table 1). Subsequently, *Neτ* increased three-fold, strongly correlating with rises in sea-level and temperature (Fig. 1A; Supplementary Fig. 3 and 6; Supplementary Table 1). Both the timing and intensity of these demographic changes concur with a three-fold expansion of bowhead whale identified over the late glacial period using ancient DNA and modelling of core suitable habitat (Supplementary Fig. 7; Ho et al. 2008; Foote et al. 2013). The end of this demographic period at 11.8 ka (Fig. 1A; Supplementary Fig. 5) matches the Pleistocene–Holocene transition at 11.6 ka, a period characterized by an ecological shift between the cold phase of the Younger Dryas and the warmer conditions of the Holocene (Walker et al. 2009). This indicates that genetic inferences can detect subtle population trends resulting from the influence of environmental changes.

**Fig. 1.**
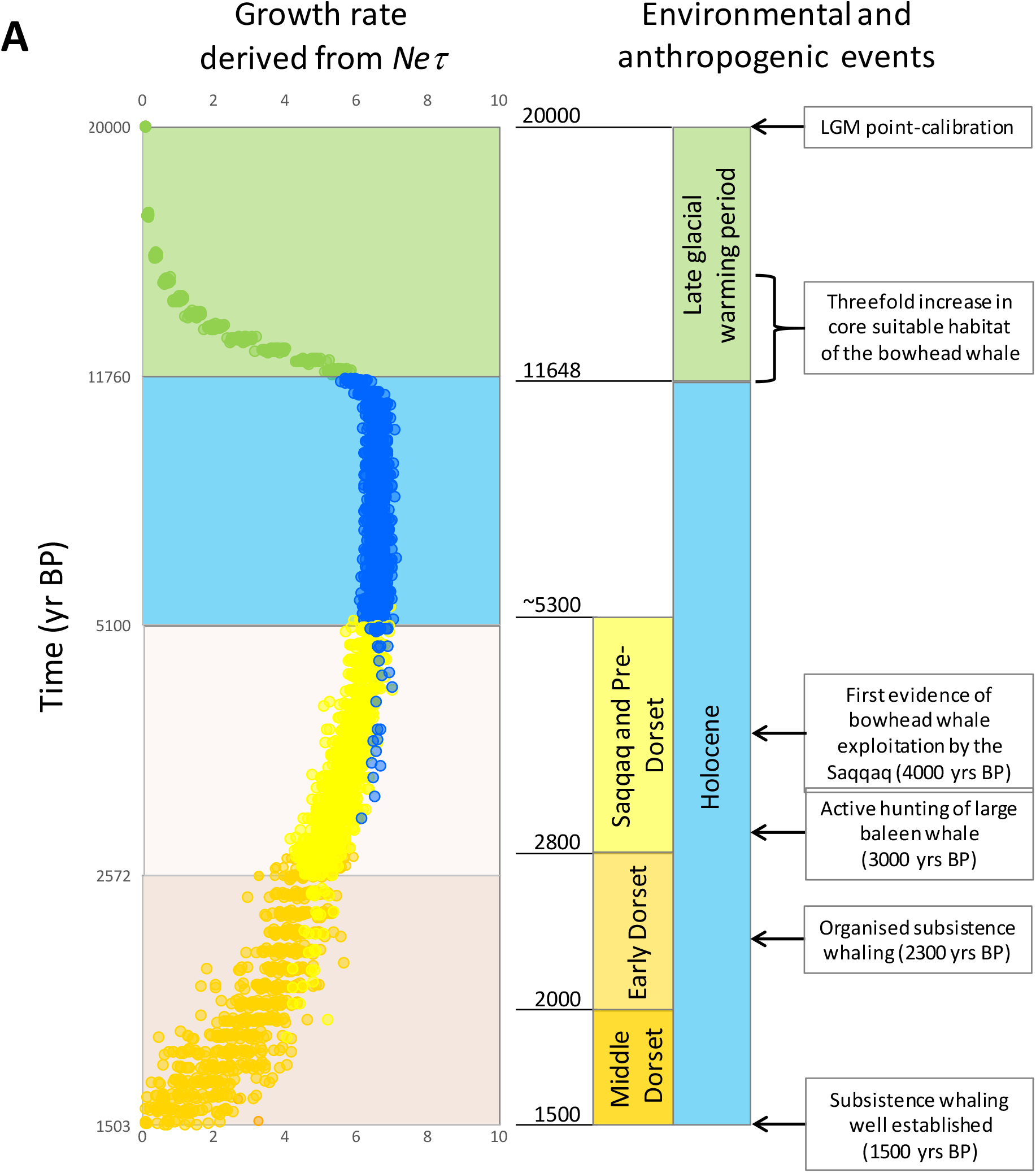
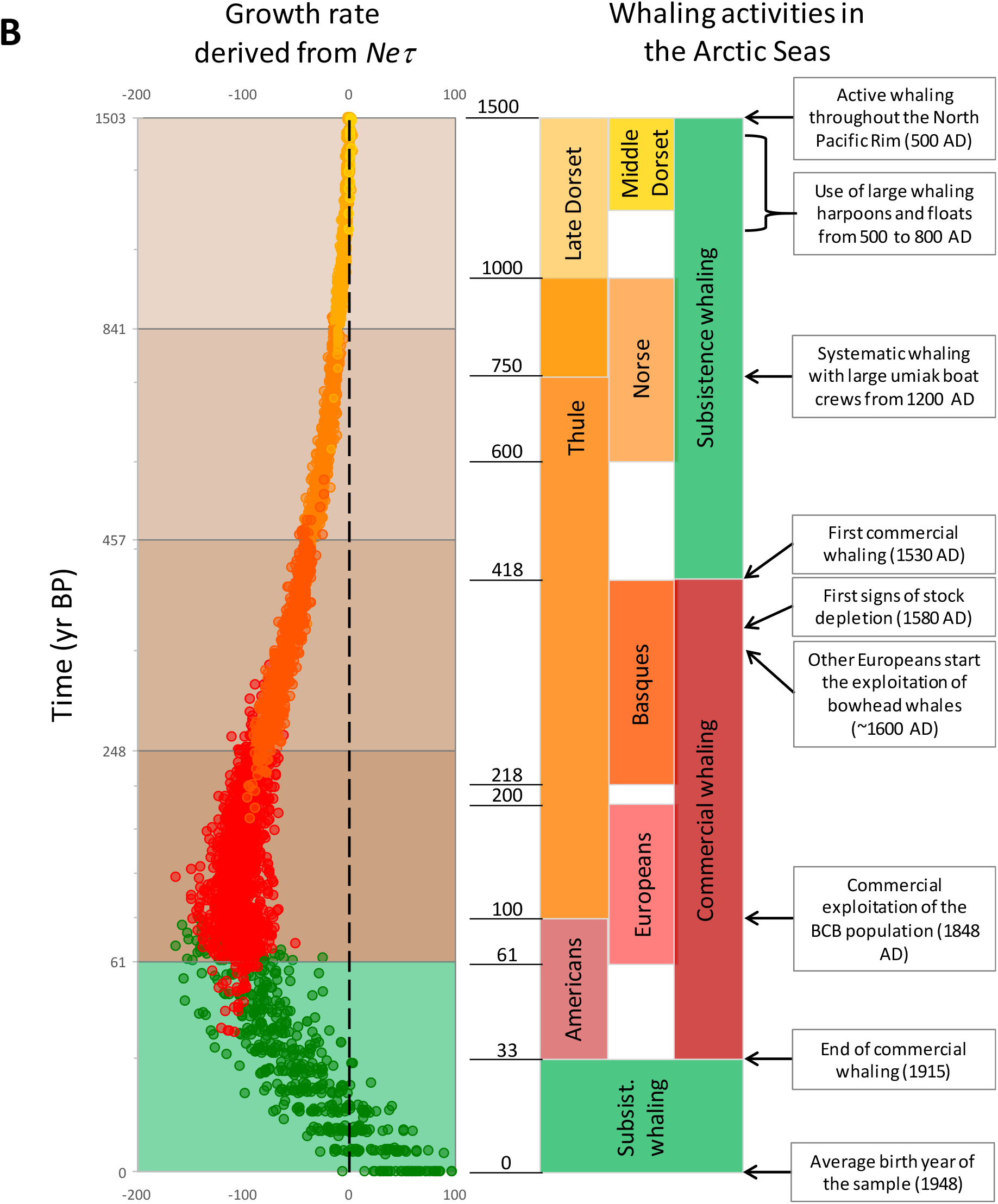
Temporal changes in the annual population growth of the bowhead whale derived from the female effective population size *(Ne)* scaled to the generation time (*τ*) and major prehistorical and historical periods in the Arctic Seas. (A) Positive population growth during the late glacial period (20.0–11.6 ka) and the Holocene (11.6–1.5 ka) with illustration of first evidence of whaling by the cultures present in the Arctic region. (B) Population growth associated with different demographic periods and significant cultures that actively hunted whales over the last 1500 years (before the mean birth year of the specimens of the sample in 1947). Different colours of population growth values in A and B represent demographic periods identified using the MCLUST algorithm (Fraley and Raftery 2003). These periods coincide with geological epochs and significant periods of active whale hunting: Late Glacial period (20.0-11.6 ka), early Holocene (11.6-5.0 ka), early evidence of whale hunting by PreDorset (5000-3000 BP), subsistence and established hunting by different phases of the Dorset culture (3000-800 BP), Thule hunting (year 1200-1500), Basque commercial exploitation (1530-1713), exploitation by other Europeans (1719-1915), the specific commercial exploitation of the BCB stock (1848-1915), and the end of commercial whaling (1915) up to the mean birth year of the specimens of the sample (1947).

The study that provided the set of reanalysed sequences (Phillips et al. 2013) identified a more ancient expansion during the glacial period (75–20 ka), but the use of a much lower substitution rate probably explains this inflation (Supplementary Fig. 2; Ho et al. 2008). Unlike the bowhead whale, most terrestrial megafauna declined in the Holartic during the late glacial (Lorenzen et al. 2011), showing that the environment is a critical driver of population fluctuations in this region. During the second demographic period identified, covering the Holocene from 11.8–5.1 ka, environmental conditions seem to have remained favourable for bowhead whales as the population growth stays constant, resulting in a 13-fold increase in *Neτ* (Fig. 1A; Supplementary Fig. 5–6; Supplementary Table 1).

### Hunting cultures are responsible for population declines in the bowhead whale

The remarkable association between species dynamics genetically-inferred here, cultural history (Seersholm et al. 2016; Savelle and Kishigami 2013) and the history of commercial whaling (Savelle and Kishigami 2013) reveals, for the first time, the impact of harvesting on the genetics of bowhead whale (Fig. 1; Supplementary Fig. 4B). From the mid-Holocene (5100-1503 BP), two demographic periods reveal declining yet positive growth values (Fig. 1A; Supplementary Fig. 4–5), which coincides with several whale-hunting cultures flourishing in the Arctic region (Raghavan et al. 2014). The Saqqaq Paleo-Inuit were the first people known to hunt bowhead whale, from around 4000 BP (Seersholm et al. 2016). The Early Dorset culture then emerged in the West Canadian Arctic region around 2800 BP (Raghavan et al. 2014), a time that coincides with the establishment of active whaling (Savelle and Kishigami 2013). Finally, the Middle Dorset culture developed the technology needed to routinely hunt large whales by 2300 BP (Raghavan et al. 2014; Seersholm et al. 2016). Declines in bowhead whale population growth could also be attributed to the species reaching carrying capacity of regional waters, linked to limited ocean productivity. At the end of these two periods at 1500 BP, *Neτ* reached its highest value at 75 430 whales which is considered here to be the pristine population size, before the first evidence of genetic decline.

The transition to the genetic decline observed from 1500 BP (Fig. 1B; Supplementary Fig. 4–5; Supplementary Table 1) coincides with the development of complex societies (Mason et al. 1998) and the establishment of organised subsistence harvesting (Savelle and Kishigami 2013) throughout the Arctic region. The subsequent demographic transitions identified correlate remarkably with the late phase of the Dorset Paleo-Inuit culture (1500–900 BP; Raghavan et al. 2014) and the rise and fall of the Thule people (900-500 BP; Savelle and Kishigami 2013). Within 400 years, harvesting for meat, oil, bones and skin by the Thule (Savelle and Kishigami 2013; Seersholm et al. 2016) is inferred to have reduced the effective size by 12%. From the sixteenth century, the Basques developed commercial whaling (Savelle and Kishigami 2013), further reducing the effective size by 17%, followed by other European (200 BP onward; Savelle and Kishigami 2013) and North Americans whalers (100 BP; Savelle and Kishigami 2013), who further induced a population decrease of 28%. By the 1930s, after the end of commercial whaling, only 38% of the original effective size remained.

### A genetic recovery is observed after the end of commercial whaling in 1915

The final demographic period identified features increasing growth rates, becoming net positive towards the present day (Fig. 1–2). I carried out simulations and confirmed that the dataset has enough resolution to detect such a recent demographic change (Supplementary Fig. 8). Based on the year collection and the average age of sampled individuals, I calculated an average birth date that can be used to calibrate the present time t0=1947 (Supplementary Fig. 9). Based on this calibration of the present, the final demographic period inferred starts around 1886. As the exploitation decreased to reach an average of 27 whales per annum from the end of commercial hunting in 1915 (Fig. 2A; Savelle and Kishigami 2013), the growth rates increased to become positive around 1934 (Supplementary Fig. 5). By 1947, *Neτ* reached 28 713 and growth reached the highest values recorded (58 whales per year; Fig. 1–2; Supplementary Table 1).

**Fig. 2.**
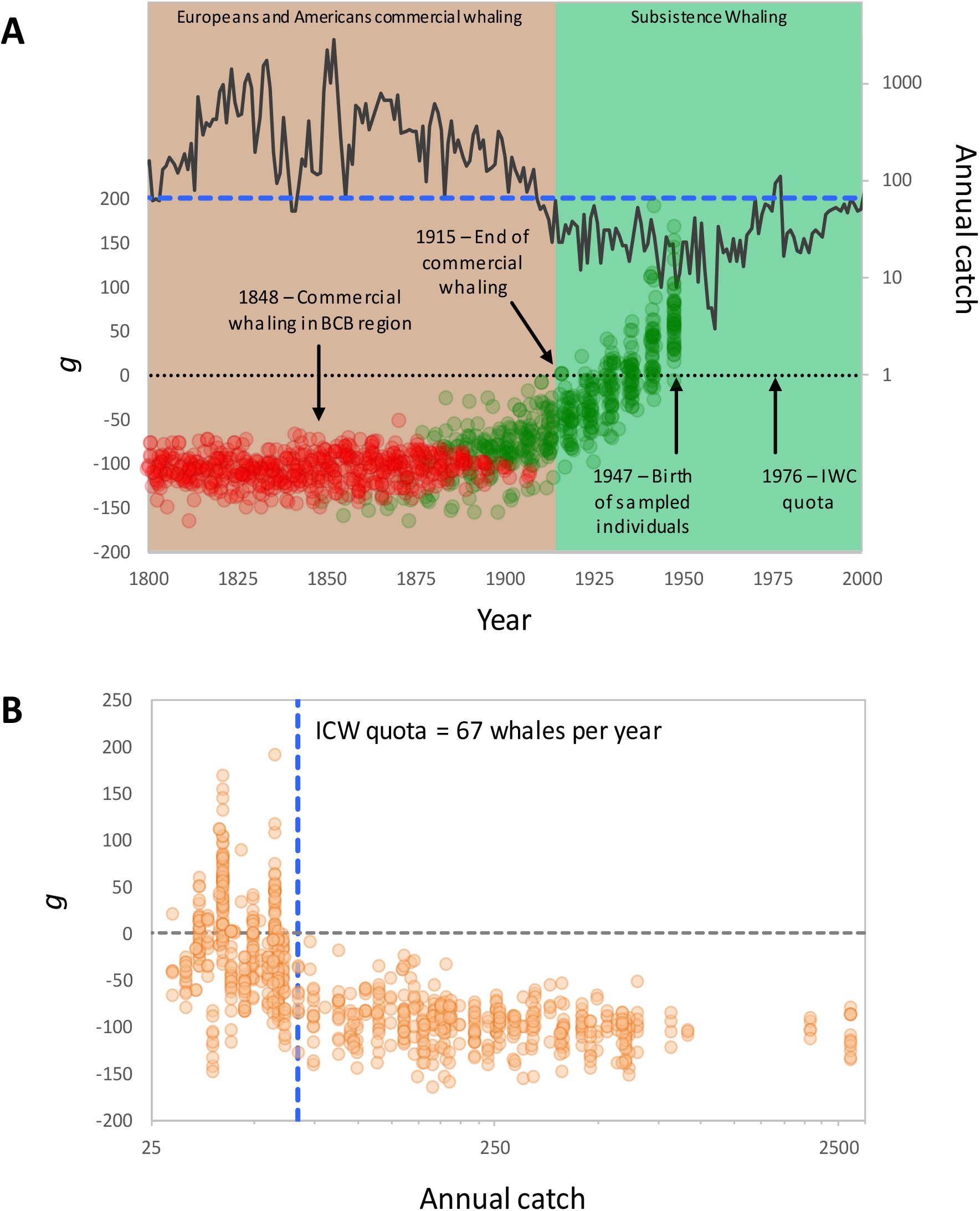
Demographic trends and recovery in the bowhead whale over the last two centuries. (A) Population growth (from 1800–1947) and annual catch in the Arctic region (1800–2000) for the bowhead whale with illustration of major events related to the exploitation. (B) Relationship between annual catch data (Brandon and Wade 2006) and annual change in effective population size (*Neτ*) of the bowhead whale from the BCB region during the period ranging from 1848–1947. The linear regression model is *y* = 2.74 + 0.08*x*, *T* = 9.62, *P* < 0.001; Spearman *Rs* = 0.3437. The annual quota established by the International Whaling Commission is illustrated by the thick blue dash lines (N = 67 bowheads) in A and B. The two colours of population growth values in A represent the demographic periods 8 and 9 identified using the MCLUST algorithm (Fraley and Raftery 2003) as in Figure 1B.

The signs of genetic recovery are concordant with an increase in abundance between 1920 and 1950 predicted by modelling studies (Witting 2013) and observed by both aboriginal people and formal surveys (George et al. 2004). When comparing annual catch and growth rates for the BCB population (Fig. 2B), genetic recovery is observed only for harvest rates lower than 67 whales per year, which is the current annual quota, established by the International Whaling Commission in 1978.

### Diverse impact of whaling on the BCB population indicates a single management unit

Heavy hunting on the North Pacific BCB population only started in 1848 with the American commercial whalers (Savelle and Kishigami 2013), yet, a clear genetic impact during the Thule and Basque periods can be detected, and these two cultures only hunted in the North Atlantic (Fig. 1B). These results indicate that the North Pacific and North Atlantic populations are demographically co-dependent, and that they respond to harvesting as a single effective population unit. This evidence for high connectivity between Pacific and Atlantic populations supports previous results inferring substantial gene flow between these predefined geographic stocks (Alter et al. 2012).

## Conclusion

Genetic inferences using a new calibration of the bowhead whale mitochondrial molecular clock have been used here to document the fine-scale temporal impact of hunting on this formerly important human resource. The approach I have taken has application to the implementation and evaluation of conservation management plans, as population recovery is observed only for annual catches comparable to the current quota. These results show that inference of population history using genetic data can be fundamentally valuable for wildlife conservation, and complementary to long-term surveys and demographic modelling studies that are often challenging (Baker and Clapham 2004; Bonebrake et al. 2010). Analyses incorporating recent samples from younger specimens should help detect demographic patterns post-1947 and confirm the recovery in more recent time.

The use of growth estimates and appropriate time calibration partially explains the detection of such a spectacular, comprehensive and sensitive genetic signatures of a dynamic population history. This remains intriguing, partly because it is based only on the median values of *Neτ* without accounting for full parameter variance, but especially since mtDNA is known to be a comparatively poor descriptor of population size in some cases (Bazin et al. 2006). These results will stimulate new studies, especially on the conservation of species that are threatened by unregulated exploitation.

## Materials and methods

### Empirical and simulated sequence datasets

I used previously published modern mtDNA sequences of the bowhead whale that include 397 bp of HVRI, 1140 bp of Cyt b, and 957 bp of ND1 (a total of 2494 bp) of 149 specimens collected in the Beaufort, Chukchi and Bering Seas (BCB) between 1995 and 2006 (Supplementary Table 2; Phillips et al. 2013). I simulated 149 non-recombining sequences of a combined 2494 nucleotides using the program FASTSIMCOAL 2 (Excoffier et al. 2013) and following two demographic scenarios. The first scenario uses the changes in *Ne* mimicking variations in sea-level observed between 135 and 0 ka to validate the molecular clock based on a first positive population growth calibrated to occur at 20 ka (Supplementary Figure 1). The second uses the changes in *Ne* recovered from the empirical data (Supplementary Figure 8) to verify that recent recovery can be detected using the genetic content observed in the empirical data (refer to the section on Extended Bayesian Skyline Plot below). The current population was set to 100 000 individuals in both cases which coincides with current estimate of the original census population size of bowhead whale with a minimum of 50 000 (Woodby and Botkin 1993, Brandon and Wade 2006). The mutation rate was set to 15.0×10^-8^ per bp per year using a HKY substitution model which is close to the rates obtained from aDNA studies on the bowhead whale (15.9× 10^-8^ per bp per year (95%HPD. 5.1-27.2×10^-8^ per bp per year); Ho et al. 2008).

### Environmental and bowhead catch data

The temperature and relative sea-level over the last 20 kyrs are environmental factors considered as potential drivers. To illustrate the temperature variation over this period, I used data derived from deuterium measurements obtained from the EPICA Dome C Ice Core from the Antarctic (Jouzel et al. 2007; http://doi.pangaea.de/10.1594/PANGAEA.683655?format=html), and applying a smoothing algorithm described in a previous study (Hoareau 2016). For the sea-level variation, I used data of relative sea-levels derived from several SPECMAP reference series that produced a standard chronology for oxygen isotope records (Waelbroek et al. 2002). To evaluate the anthropogenic effect on the bowhead whale population, I use the annual catch data obtained for the BCB population specifically (Brandon and Wade 2006) and for the overall species (Higdon, 2010).

### Inference of past population dynamics using the extended Bayesian Skyline Plot model

To infer the population history, I first applied the extended Bayesian Skyline plot model (EBSP; Heled and Drummond 2008) to the data to obtain the variation in *Neτ* (with *τ* the generation time) over the time parameter *t*. This model is available in the program BEAST version 1.7.2 (Drummond and Rambaut 2012). I selected the HKY model of nucleotide substitution based on the Bayesian information criterion available in MEGA 6.06 (Tamura et al. 2013) which also matches finding in another study (Foote et al. 2013). At first, the molecular clock was left uncalibrated so that the posterior distribution of both the effective population size (*Neτ*) and the associated time parameter (*t*) are scaled to the rate of changes per site (not calibrated according to calendar times). The MCMC parameters were set to 110 million iterations and, after discarding 10% of samples as burn-in, 10 000 genealogies and model parameters remained for further analyses. To ensure the quality of the runs, I used TRACER v1.6 (http://tree.bio.ed.ac.uk/software/tracer) to verify that the effective sample sizes (ESS) of all parameters were above 200 as suggested by the BEAST guidelines. Several hundreds of independent runs of the empirical dataset were done to ensure the recovery of 46 convergent runs. These were used as independent records of the past population dynamics of the bowhead whale that provided a variance estimates of time and demographic parameters.

### Calculating the population growth rate

To provide more informative parameters of the demographic trajectories, I derived the annual population growth using the median values of *Neτ* and *t* recovered for each of the convergent runs using the following equation [*R* = d*Neτ* /d*t* = (*Neτ*_n_ – *Neτ*_n+1_)/(*t*_n+1_ – *t*_n_)] (with *Ne*: effective population size, *τ* generation time of the species, *t*: time and n: number of coalescent events back in time). This provided the rate estimates of the population growth at every point in time associated with the coalescent events recorded in the given dataset (n = 148 coalescent events for a sample size of 149), and this back to the most recent common ancestor of the sequence dataset.

### Calibration of molecular rates

The calibration relies on the single assumption that the bowhead whale has a first positive population growth rate (*R* > 0) at 20 ka (Fig. 1A). This assumption is based on the previous palaeoecological results obtained on bowhead whale that show that the habitat was minimal during the LGM, and then increased to become maximal during the Holocene (Foote et al. 2013). The mutation rate obtained using this simple calibration was then used to calibrate the EBSP analyses, which comprise the time parameters associated with the population growth rate estimates (Fig. 1A).

Because the bowhead is long live, I used an average birth year of 1947 (supplementary Fig. 9) as a proxy for the present time (t = 0) of the genetic sample. Birth year was calculated by subtracting age estimate and collection year of 50 specimens analysed by George and co-workers (2011). Some specimens used in the present work were also part of this previous study. Here, I assume that the specimens analysed have the same birth year distribution as the whole sample analysed by George and co-authors (2011). As the year 1947 is very close to 1950, which is routinely defined as the “present” in studies using radiocarbon dating, I express dates in years Before Present (BP) for all values ranging from around 5000 to 2000 BP. All dates that are more recent than 2000 BP are expressed in calendar years.

### Growth trends and transition time estimates

To explore the possibility of distinct demographic trajectories (e.g. growth trends) over time, I applied a clustering algorithm on the pool of 46 full datasets (with good ESSs values) of population growth and time estimates (6808 values for each parameter). This MCLUST algorithm available in R (Fraley and Raftery 2003) uses a Bayesian approach to minimize the variance within group. The recovered clusters correspond to distinct time periods with different average growth estimates. I calculated the transition dates (and 95% percentile) between these periods using a logistic regression approach available in the R package aod (Analysis of Overdispersed Data version 1.3). I then compared these transition dates to periods that are known to influence the demography of the bowhead whale (environmental changes or human activities) and obtained from archaeological data as well as historical records. Scatter plots with linear fit and regression models between the different parameters were performed using the R statistical framework.

## Supporting information

Supplementary Table 2 - Sequence information

## Supplementary Material

Supplementary data are available as a separate file online.

## Acknowledgements

This work was supported by the Research and Innovation Support of University of Pretoria, and the author benefitted from the University of Pretoria’s research fellowship programme. This work has benefitted from the technical support and the bioinformatic resources of the Centre for Bioinformatics and Computational Biology of the University of Pretoria and the Centre for High Performance Computing (CHPC) from the South African Department of Science and Technology (DST). The author benefitted from insights from Profs Rasmus Nielsen and Alexei Drummond on the simulations. The author is grateful to Prof Fourie Joubert and Dane Kennedy for precious bioinformatic support, and Prof Michael Bruford, Dr Michael Cunningham and Mrs Hanlie Griesel for providing constructive comments on earlier drafts of the manuscript.

