## Supplementary Table 1 and Figures 1-9 for "Whales and Men: genetic inferences uncover a detailed history of hunting in bowhead whale"

**Supplementary Table 1.** Details of the estimates of transition times (as defined in Supplementary Figure S5), and associated $Ne\tau$ (Female effective population size scaled to generation times) and *g* (intrinsic growth rate derived from *Ne*) estimates.

| Time (95% percentile) | Year AD | $Ne\tau$ (95% percentile) | *g* (95% percentile) |
|---|---|---|---|
| 0 | 1947 | 28713 (24113-31535) | 58.7 (15.4–152.5) |
| 13 (0-30) | 1934 | 28257 (23320-31481) | -3.5 (-79.2–110.8) |
| 32 (12-52) | 1915 | 28643 (23930-31654) | -52.8 (-118.9–20.0) |
| 61 (33-88) | 1886 | 30729 (26446-34397) | -94.4 (-142.2–-33.6) |
| 248 (197-299) | 1699 | 49510 (44991-54239) | -86.5 (-111.0–-66.1) |
| 457 (403-511) | 1490 | 62665 (59862-65889) | -40.2 (-52.4–29.3) |
| 841 (752-930) | 1106 | 72039 (70166-74421) | -12.3 (-18.5–-7.6) |
| 1503 (1337-1669) | 444 | 75430 (74180-76864) | 0.1 (-3.0–-2.6) |
| 2572 (2160-2984) | NA | 72516 (70105-74656) | 4.6 (3.3–5.6) |
| 5100 (4317-5882) | NA | 58374 (53451-63039) | 6.4 (5.8–6.8) |
| 11760 (11740-11780) | NA | 14608 (13266-15982) | 5.6 (5.2–6.2) |
| 20000 | NA | 4510 (4391-4661) | 0.1 (0.1–0.2) |

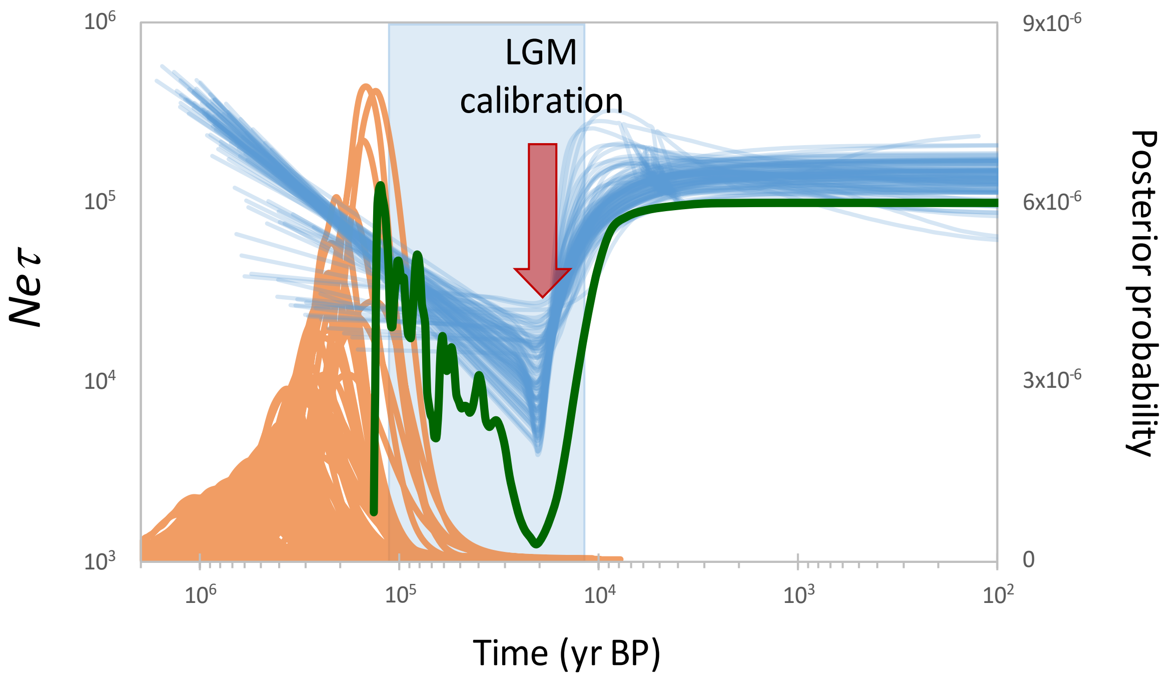


**Fig. S1.** Validation of the LGM-point calibration method using a simulation study. The *x*-axis is in calendar years before present; the *y*-axis is the product of female effective population size and generation time ($Ne\tau$). Population size reconstructions (blue curves) and posterior distribution of the Time to the Most Recent Common Ancestor (orange curves) for 99 datasets simulated following a demographic trajectory mimicking sea-level changes over the last 135 ka BP (green curves) and using the extended Bayesian Skyline plot model as well as a calibration assuming a bottleneck during the LGM with a first positive population growth around 20 ka. The last glacial period is indicated by the light shading in the background.

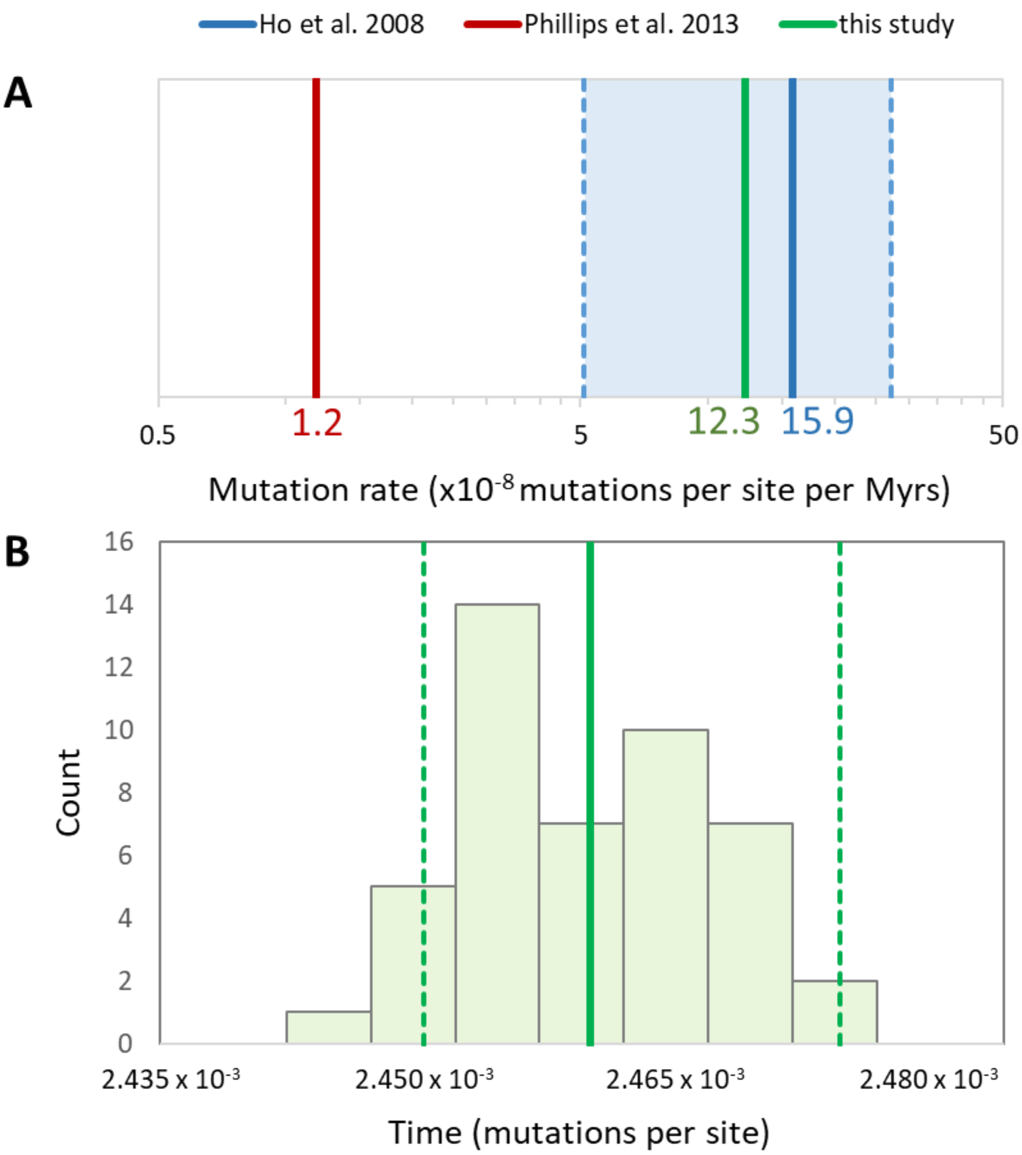


**Fig. S2.** Mutation rates used to calibrate the demography of bowhead whales (A) and timing of the earliest positive population growth rate measured in units of mutations per site in the present study (B). In A, mutation rates resulting from different calibration methods are illustrated including the LGM-point calibration (green line; present study), a calibration based on ancient DNA (thick and dashed blue lines; Ho et al. 2008) and a phylogenetic calibration (red line; Phillips et al. 2013) are illustrated. LGM-point calibration assumes that the first coalescent event characterized by a positive growth occurs at 20 ka. The mean mutation rate is $1.230 \times 10^{-7}$ (95% percentile: $1.225 \times 10^{-7}$–$1.238 \times 10^{-7}$) change per site per million years and the mean time considered for calibration is $2.460 \times 10^{-3}$ (95% percentile: $2.450 \times 10^{-3}$–$2.475 \times 10^{-3}$) in mutation per site.

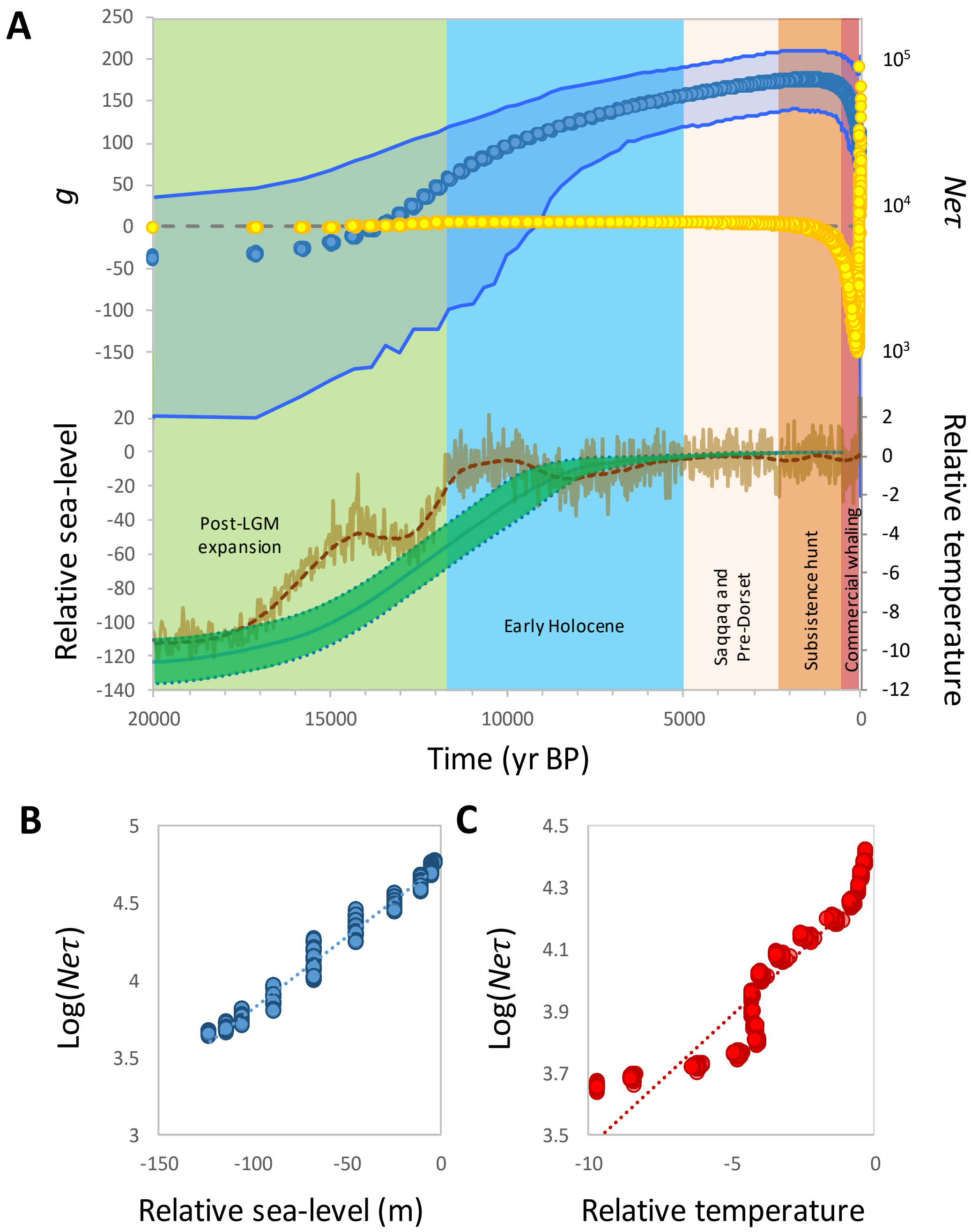


**Fig. S3.** Association between the demography of the bowhead whale and environmental factors. (A) Female effective population size (*Ne*$\tau$; blue dots) and derived population growth rate (*g*; orange dots) of bowhead whales from the Bering-Chukchi-Beaufort region and variation in relative sea-level (Waelbroeck et al. 2002) and relative temperature (Jouzel et al. 2007) over the last 20 kyrs. The periods known to play a role in bowhead demography are illustrated in the background. (B) Scatter plot with linear fit showing female effective population size ($Ne\tau$) versus relative sea-level between 5–20 ka. The regression model is $y = 4.7471 + 0.0093x$, slope test $t = 297.36$, $P < 0.0001$; Pearson $R = 0.9905$, $F = 8.84 \times 10^4$, $P < 0.0001$. (C) Scatter plot with linear fit showing female effective population size ($Ne\tau$) versus relative temperature between 10–20 ka. The regression model is $y = 4.3144 + 0.0854x$, slope test $t = 70.88$, $P < 0.0001$; Pearson $R = 0.9313$, $F = 5024$, $P < 0.0001$.

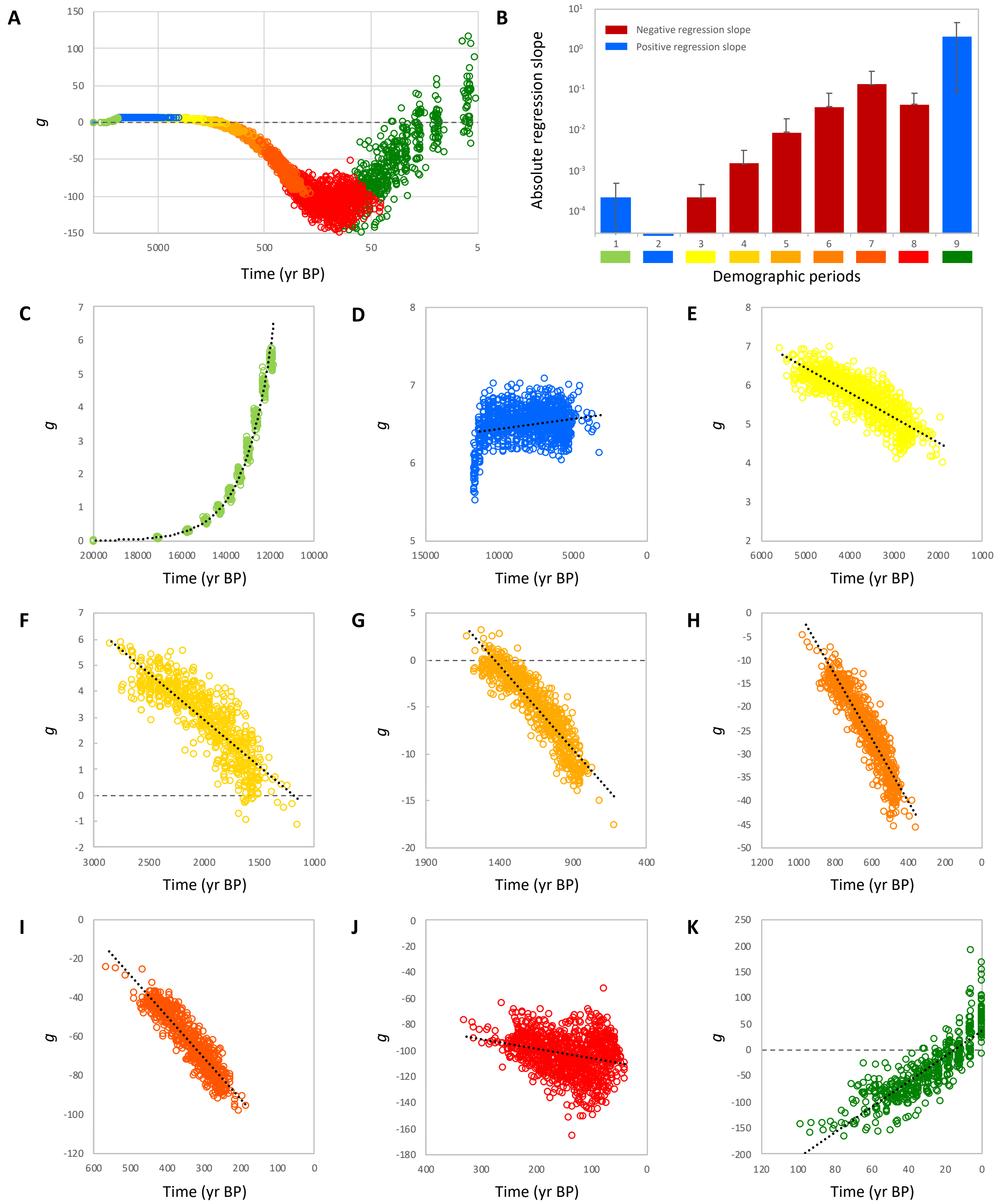


**Fig. S4.** Population growth rate (g) derived from the female effective population sizes of the bowhead whale in relation to calibrated time for the nine demographic periods recovered from the Bayesian clustering analysis. (A) Overview of the nine demographic periods illustrating the variation in population growth rate over time between 20 ka and the present ($t$ = 0). (B) Regression slopes assuming a linear model between population growth rate and time for the different demographic periods recovered. (C) Demographic period 1 with a relation matching an exponential model: $y = 68150 \times e(-8\times10^{-4}x)$; $R = 0.9981$, $F = 1.344\times10^{5}$, $P < 0.0001$. (D) Demographic period 2 with a relation matching a linear regression model $y = 6.6853 - 2.44\times10^{-5}x$; Pearson $R = 0.2272$, $F = 66.84$, $P < 0.0001$. (E) Demographic period 3 with a relation matching a linear regression model $y = 3.2635 + 0.0006x$; Pearson $R = 0.8373$, $F = 2287$, $P < 0.0001$. (F) Demographic period 4 with a relation matching a linear regression model $y = -4.2989 + 0.0036x$; Pearson $R = 0.8565$, $F = 1895$, $P < 0.0001$. (G) Demographic period 5 with a relation matching a linear regression model $y = -25.728 + 0.018x$; Pearson $R = 0.9121$, $F = 3251$, $P < 0.0001$. (H) Demographic period 6 with a relation matching a linear regression model $y = -67.107 + 0.0673x$; Pearson $R = 0.9196$, $F = 3517$, $P < 0.0001$. (I) Demographic period 7 with a relation matching a linear regression model $y = -134.26 + 0.211x$; Pearson $R = 0.9228$, $F = 3585$, $P < 0.0001$. (J) Demographic period 8 with a relation matching a linear regression model $y = -113.8 + 0.0749x$; Pearson $R = 0.2792$, $F = 88.19$, $P < 0.0001$. (K) Demographic period 9 with a relation matching a linear regression model $y = 36.55 – 2.4393x$; Pearson $R = 0.8433$, $F = 1211$, $P < 0.0001$.

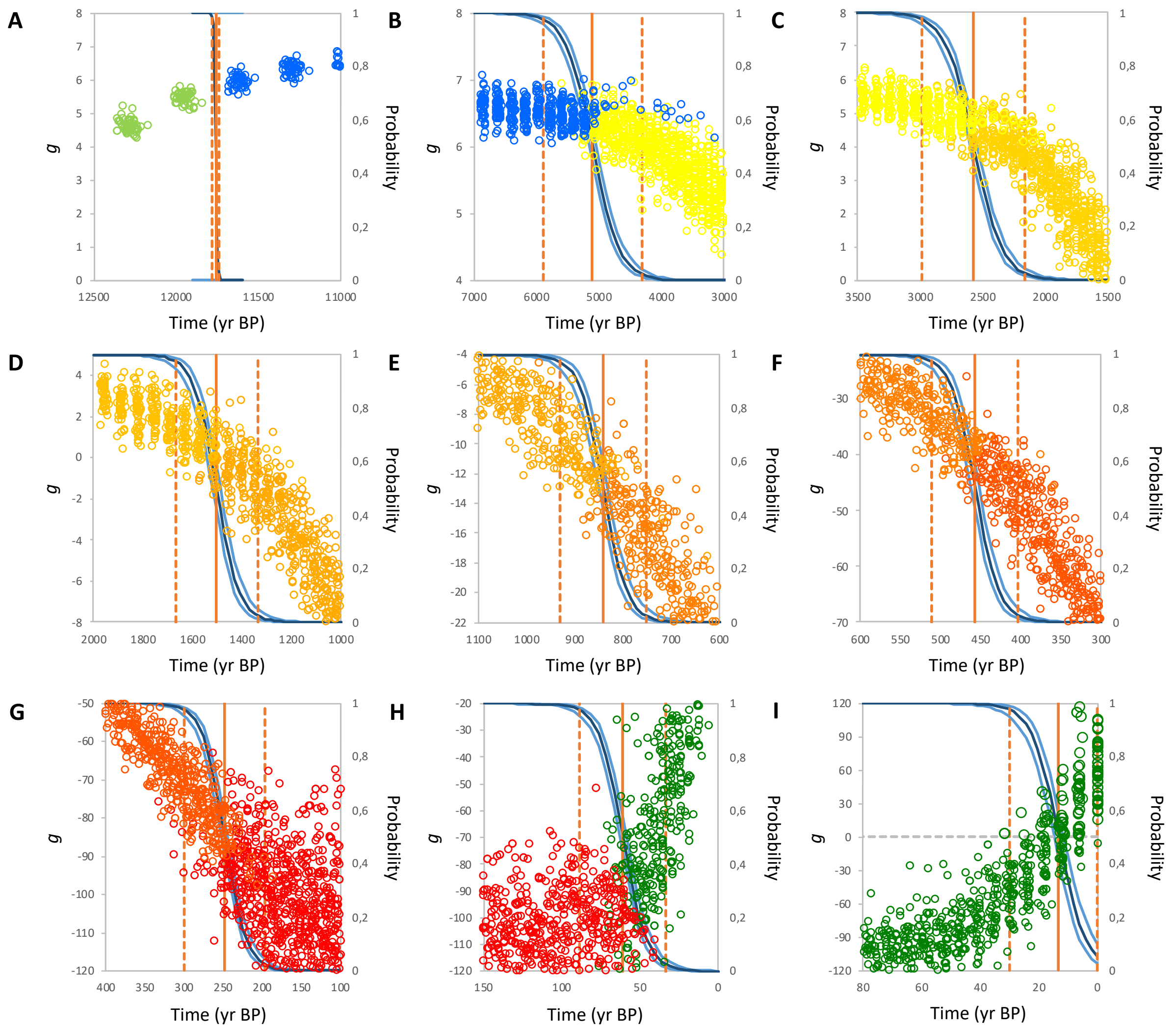


**Fig. S5.** Transition time between successive demographic periods of the bowhead whale estimated using the curve of a logistic regression. In each case, the growth values of the most recent demographic period are assigned the value 0, and the previous period the value 1. The logistic regression curves (blue) relate the predicted probability of transition of the two demographic periods (*y*-axis) over time (*x*-axis in years before present). The vertical continuous lines indicate the time at which the two demographic periods are equally likely (0.5), and the dash lines indicate the 95% percentile of the transition between periods (0.975 and 0.025 respectively). (A) Transition time between the demographic periods 1 and 2: $t_A$ = 11760 (95 percentile: 11740–11780) year before present (BP); (B) Transition between the periods 2 and 3: $t_B$ = 5100 (4317–5882) BP; (C) Transition between the periods 3 and 4: $t_C$ = 2572 (2160–2984) BP; (D) Transition between the periods 4 and 5: $t_D$ = 1503 (1337–1669) BP; (E) Transition between the periods 5 and 6: $t_E$ = 841 (752–930) BP; (F) Transition between the periods 6 and 7: $t_F$ = 457 (403–511) BP; (G) Transition between the periods 7 and 8: $t_G$ = 248 (197–299) BP; (H) Transition between the periods 8 and 9: $t_H$ = 61 (33–88) BP; (I) Transition between the periods 9 (negative growth values) and 9 (positive growth values): $t_I$ = 13 (0–30) BP.

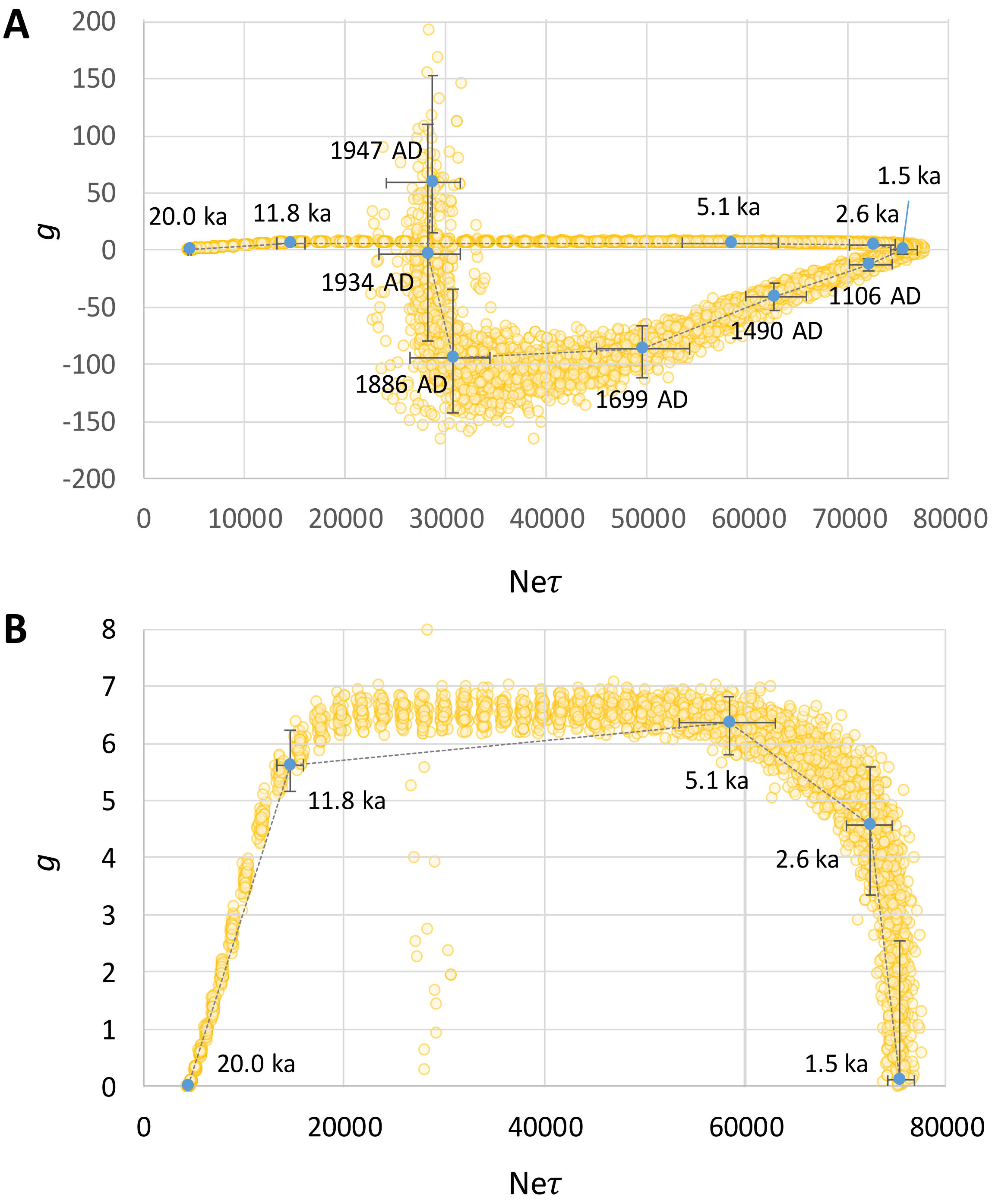


**Fig. S6.** Intrinsic growth rate ($g$) as a function of the female effective population size (Ne$\tau$), for the BCB population of bowhead whales over 20 ka. (A) Overall relationship. (B) Details of the relationship between 20 ka and 1500 BP, while the growth is still positive. The median values (95% percentile) of the intrinsic growth are indicated for the different demographic transitions detected over the 20 krys history. The estimates of Female Effective Population Size (*Ne*) are scaled to the generation times ($\tau$).

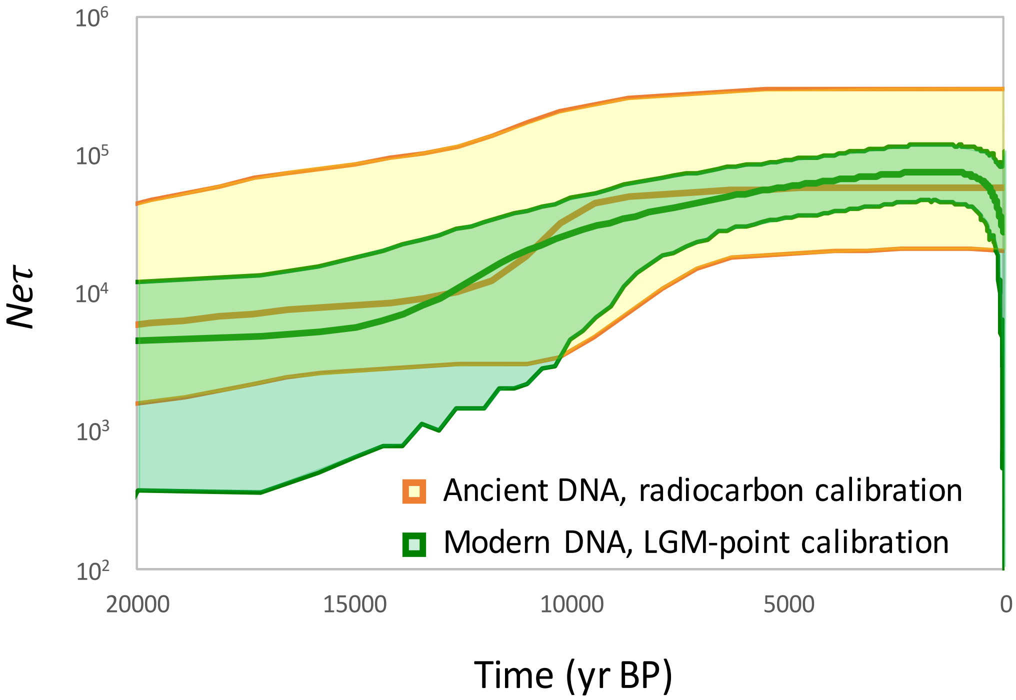


**Fig. S7.** Comparison of demographic reconstructions based on ancient DNA (Ho et al. 2008) and the LGM-point calibration based on demographic assumption. Time is expressed in years before present (BP) and the female effective population size is scaled to the generation time ($Ne\tau$). The mutation rate calibrated from the modern sequences is $12.3\times10^{-8}$ per bp per year and the mutation rate obtained using ancient DNA is $15.9\times10^{-8}$ per bp per year (Ho et al. 2008).

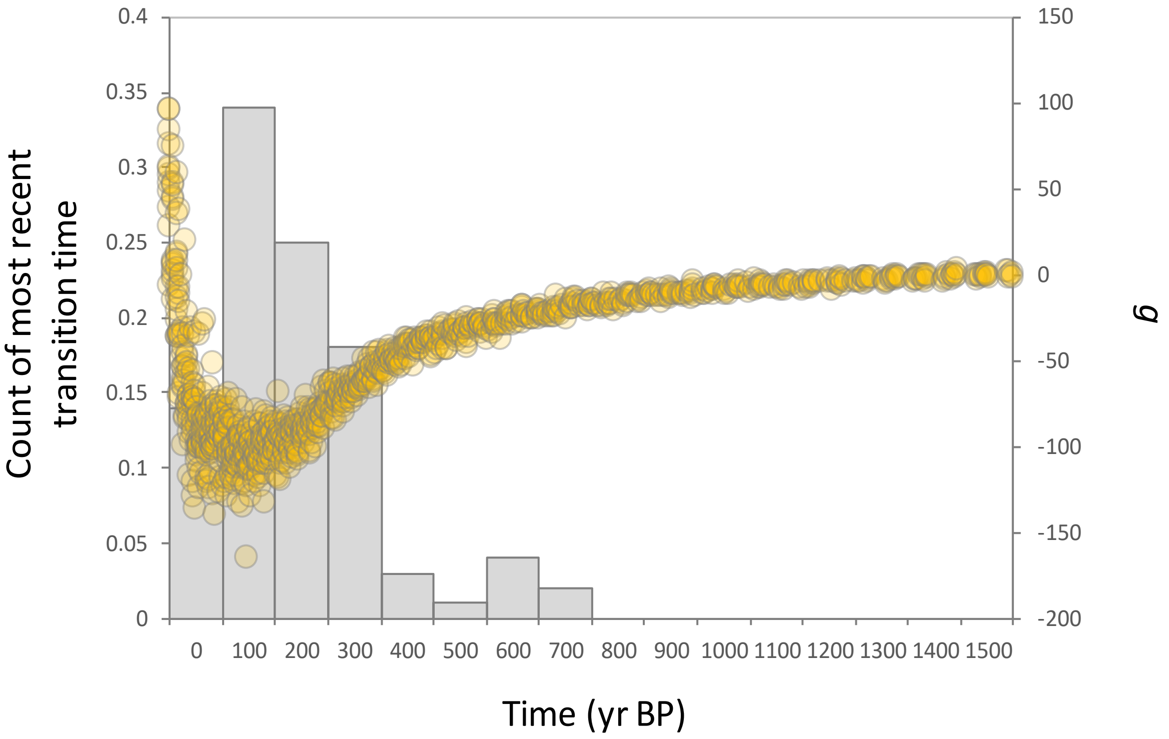


**Fig. S8.** Transition times for simulated demographic recoveries (bar plot) and variation of intrinsic population growth for the bowhead whale (circles). Most estimates of the transition time are found in the vicinity of the change observed in the bowhead whale (mode = 100 years, median = 157 years, mean = 195, 95% percentile: 23-620 years).

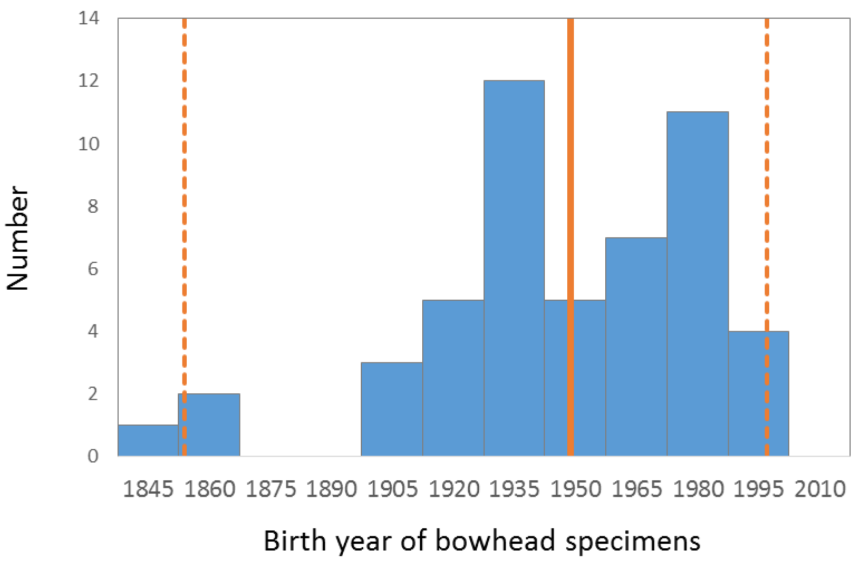


**Fig. S9.** Distribution of birth year for specimens of bowhead whale spanning the range of collection dates of the current sample. The mean age of 46 years old ranging from 6 to 149 years old were obtained from previously published data (N=50, George et al. 2011). Subtracting the age to the mean capture date, the mean birth year is 1947 (95% percentile: 1860–1991) which is used as a calibration for t = 0 of the genetic sample.
